# Mechanism of RNA recognition by a Musashi RNA-binding protein

**DOI:** 10.1101/2020.10.30.362756

**Authors:** Jinan Wang, Lan Lan, Xiaoqing Wu, Liang Xu, Yinglong Miao

## Abstract

The Musashi RNA-binding proteins (RBPs) regulate translation of target mRNAs and maintenance of cell stemness and tumorigenesis. Musashi-1 (MSI1), long considered as an intestinal and neural stem cell marker, has been more recently found to be overexpressed in many cancers. It has served as an important drug target for treating acute myeloid leukemia and solid tumors such as ovarian, colorectal and bladder cancer. One of the reported binding targets of MSI1 is Numb, a negative regulator of the Notch signaling. However, the dynamic mechanism of Numb RNA binding to MSI1 remains unknown, largely hindering effective drug design targeting this critical interaction. Here, we have performed all-atom simulations using a robust Gaussian accelerated molecular dynamics (GaMD) method, which successfully captured spontaneous and highly accurate binding of the Numb RNA from bulk solvent to the MSI1 protein target site. GaMD simulations revealed that Numb binding to MSI1 involved largely induced fit in both the RNA and protein. The simulations also identified important low-energy intermediate conformational states during RNA binding, in which Numb interacted mainly with the β2-β3 loop and C terminus of MSI1. The mechanistic understanding of RNA binding obtained from our GaMD simulations is expected to facilitate rational structure-based drug design targeting MSI1 and other RBPs.

## Introduction

Protein-RNA interactions play crucial roles in various cellular activities and their dysfunction leads to a wide range of human diseases^1–3^. Identification of small molecules that modulate interactions between RNA-binding proteins (RBPs) and RNA is progressing rapidly. It represents a novel strategy for discovery of drugs with new mechanisms^1^. The Musashi (MSI) RBPs have been shown to regulate translation of target mRNAs and participate in the maintenance of cell stemness and tumorigenesis. They have been suggested as potential drug targets for treating many types of human cancer, including acute myeloid leukemia, ovarian cancer, colorectal cancer and bladder cancer^4^. The MSI protein family has two members: MSI1 and MSI2. Each MSI protein contains two N-terminal RNA recognition motifs (RRM1 and RRM2) that mediate the binding to their target mRNAs^5^. MSI1 binds to the 3’-untranslated region of Numb mRNA and represses its translation, which confers to the upregulation of Notch signaling. This leads to increased cell proliferation and survival, and decreased apoptosis of cancer cells^4^. Understanding the molecular mechanism of MSI1-Numb RNA interaction is important in both basic biology and applied medical research.

Rational design of small molecules targeting protein-RNA interactions requires structural characterizations of the RBP-RNA complexes. Due to high flexibility of MSI proteins and the lack of potent ligands, only a few MSI structures have been resolved so far, including the *apo* structure of MSI1/2-RRM1^6–10^ and RNA-bound structure of MSI1^11–12^. These structures have greatly facilitated structure-based modeling and drug design targeting the MSI-RNA interactions^13–16^. For example, we have recently identified one compound Aza-9 by combining fluorescence polarization (FP) assay, surface plasmon resonance (SPR), nuclear magnetic resonance (NMR) spectroscopy and molecular docking^16^. However, the experimental structures are rather static images and the dynamic mechanism of MSI-RNA interactions remains unknown, which has largely hindered the development of potent inhibitors of MSI proteins.

Molecular dynamics (MD) is a powerful technique that enables all-atom simulations of biomolecules. MD simulations are able to fully account for the flexibility of the RBP and RNA during their interactions^17–19^. In 2015, Krepl et al.^20^ provided systematic benchmarking data by simulating six structurally diverse protein/RNA complexes over multiple microsecond timescale MD runs and evaluating the simulations’ stability. Their results suggested that the current force fields are able to handle microsecond MD simulations of protein/RNA complexes in many cases. For most systems, MD was possible to achieve a good but imperfect agreement with the experimental structure. However, MD could not maintain the initial experimental structure in one among six cases (3K5Y). The same group further presented a joint MD and NMR study to interpret and expand the available structural data of two RBPs bound with their single-stranded target RNAs^21^. They collected more than 50 μs simulations and showed that the MD simulation was robust enough to reliably describe structural dynamics of RBP-RNA complexes.^21^ However, due to the slow dynamics and limited simulation timescale, it is rather challenging for conventional MD (cMD) simulations to sufficiently sample RBP-RNA interactions and obtain proper free energy profiles to quantitatively characterize RBP-RNA interactions.

To overcome limitations of cMD, enhanced sampling methods have been developed to improve biomolecular simulations^22–28^. Enhanced sampling methods have also been applied in studies of RBP-RNA interactions, including the steered MD^29^, umbrella sampling^29^ and metadynamics^30–32^. Nevertheless, these enhanced sampling methods require predefined collective variables and may introduce constrains on the conformational space of the proteins. In this context, Gaussian accelerated MD (GaMD) has been developed to allow for unconstrained enhanced sampling and free energy calculations of large biomolecules.^28, 33–34^ GaMD has been applied to successfully simulate protein folding and conformational changes^33–38^, ligand binding^33–35, 39–44^, protein-protein/membrane/nucleic acid interactions^45–50^. More recently, GaMD simulations have successfully captured spontaneous binding of RNA to the human respiratory syncytial virus M2-1 protein^51^.

In this study, we have performed all-atom enhanced sampling simulations using GaMD on MSI1-Numb RNA interactions started from the NMR structure in the bound state^11^, as well as the unbound state with the Numb RNA moved far away from the MSI1 protein surface (**Table 1**). While the NMR structure was found to maintain the bound conformation of MSI1-Numb during the GaMD simulations, further simulations captured complete binding of the Numb RNA to the MSI1 protein. The simulations thus allowed us to characterize structural flexibility and free energy landscapes of the MSI1-Numb RNA interactions, which provided important insights into the mechanism of RNA recognition by the MSI1 RBP.

**Table 1.** Summary of GaMD simulations performed on the MSI1 protein with RNA Numb started from the Bound and Unbound states.

| **System** | **$N_{atoms}$** | **Length(ns)** | **Boost potential ( kcal/mol)** |
| --- | --- | --- | --- |
| MSI1-Numb (Bound) | 29,368 | 500 x 3 | $14.08 \pm 3.79$ |
| MSI1-Numb (Unbound) | 29,569 | 1200 x 4 | $11.53 \pm 3.36$ |

## Methods

### Gaussian accelerated molecular dynamics (GaMD)

GaMD enhances the conformational sampling of biomolecules by adding a harmonic boost potential to reduce the system energy barriers.^35^ When the system potential 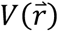 is lower than a reference energy E, the modified potential 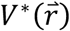 of the system is calculated as:

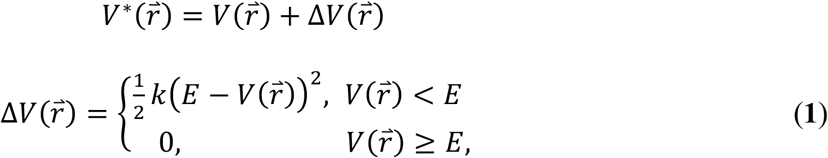

Where k is the harmonic force constant. The two adjustable parameters E and k are automatically determined on three enhanced sampling principles. First, for any two arbitrary potential values 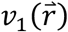 and 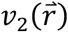 found on the original energy surface, if 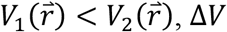, Δ*V* should be a monotonic function that does not change the relative order of the biased potential values; i.e., 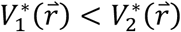. Second, if 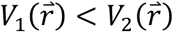, the potential difference observed on the smoothened energy surface should be smaller than that of the original; i.e., 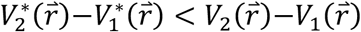. By combining the first two criteria and plugging in the formula of 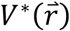 and Δ*V*, we obtain

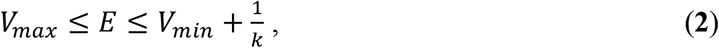

Where *V*_*min*_ and *V*_*max*_ are the system minimum and maximum potential energies. To ensure that **Eq. 2** is valid, *k* has to satisfy: *k* ≤ 1/(*V*_*max*_ − *V*_*min*_. Let us define: *k* = *k*_0_ ∙ 1/(*V*_*max*_ − *V*_*min*_, then 0 < *k*_0_ ≤ 1. Third, the standard deviation (SD) of Δ*V* needs to be small enough (i.e. narrow distribution) to ensure accurate reweighting using cumulant expansion to the second order: σ_Δ*V*_ = *k*(*E* − *V*_*avg*_)σ_*V*_ ≤ σ_0_, where *V*_*avg*_ and σ_*V*_ are the average and SD of Δ*V* with σ_0_ as a user-specified upper limit (e.g., 10*k*_*B*_*T*) for accurate reweighting. When E is set to the lower bound *E* = *V*_*max*_ according to **Eq. 2**, *k*_0_ can be calculated as

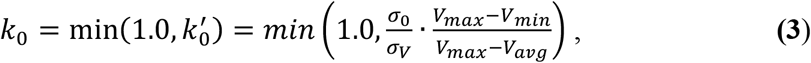

Alternatively, when the threshold energy E is set to its upper bound *E* = *V*_*min*_ + 1/*k*, *k*_0_ is set to:

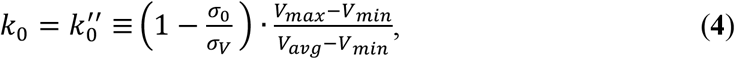

If 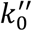 is calculated between 0 and 1. Otherwise, *k*_0_is calculated using **Eq. 3**.

### Energetic Reweighting of GaMD Simulations

For energetic reweighting of GaMD simulations to calculate potential of mean force (PMF), the probability distribution along a reaction coordinate is written as *p*\*(*A*). Given the boost potential Δ*V*(*r*) of each frame, *p*\*(*A*) can be reweighted to recover the canonical ensemble distribution *p*(*A*), as:

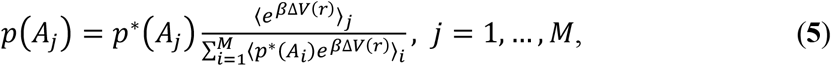

where *M* is the number of bins, *β* = *k*_*B*_*T* and 〈*e*^*β*Δ*V*(*r*)^〉_*j*_ is the ensemble-averaged Boltzmann factor of Δ*V*(*r*) for simulation frames found in the *j*^th^ bin. The ensemble-averaged reweighting factor can be approximated using cumulant expansion:

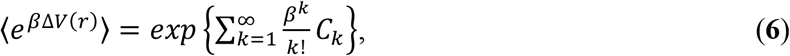

where the first two cumulants are given by:

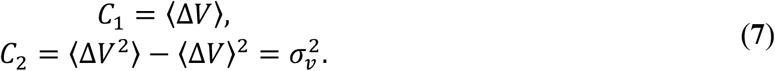

The boost potential obtained from GaMD simulations usually follows near-Gaussian distribution ^34^. Cumulant expansion to the second order thus provides a good approximation for computing the reweighting factor ^35, 52^. The reweighted free energy *F*(*A*) = −*k*_*B*_*T* ln *p*(*A*) is calculated as:

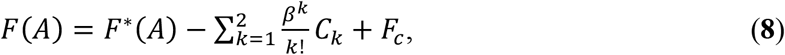

where *F*\*(*A*) = −*k*_*B*_*T* ln *p*\*(*A*) is the modified free energy obtained from GaMD simulation and *F*_*c*_ is a constant.

### System setup

Two models were prepared for simulations of MSI1-RNA interactions. One was obtained from the first model of NMR structure of the Numb RNA-bound MSI1 protein (PDB: 2RS2^11^, denoted as the “Bound” state). Another one was generated with the first model by moving the Numb RNA ~30 Å away from its binding site in MSI1 (denoted as the “Unbound” state). The two systems were solvated in explicit water using *tleap* in the AMBER 20 package^53^. The system charges were then neutralized at 0.15M NaCl using *tleap*. The AMBER ff14SBonlysc force field parameters^54^, RNA.LJbb^55^ and TIP3P model^56^ were applied for the protein, RNA and water molecules, respectively. Each system was minimized using steepest descent for 50,000 steps and conjugate gradient for another 50,000 steps. After minimization, the system was heated from 0 to 300 K in 1 ns simulation by applying 1 kcal/(mol•Å^2^) harmonic position restraints to the protein and RNA heavy atoms with a constant number, volume and temperature (NVT) ensemble. Each system was further equilibrated using a constant number, pressure and temperature (NPT) ensemble at 1 atm and 300 K for 1ns with same restraints as in the NVT run. Another 1.2 ns cMD simulations were performed to collect potential energy statistics (including the maximum, minimum, average and standard deviation). Then 24 ns GaMD equilibration after applying the boost potential was performed. Finally, three independent 500 *ns* and four independent 1,200 ns GaMD production runs with randomized initial atomic velocities were performed on the bound and unbound MSI1-Numb systems, respectively (**Table 1**). Simulation frames were saved every 0.4 ps for analysis.

### Simulation Analysis

CPPTRAJ^57^ and VMD^58^ were used to analyze the GaMD simulations. Important reaction coordinates were identified from the simulation trajectories such that they involved dynamic regions (e.g., the β2-β3 loop of MSI1) and could be used to differentiate conformational states of the MSI1-Numb system. Therefore, root-mean-square deviations (RMSDs) of the backbone of core RNA (central three nucleotides UAG in Numb) and the β2-β3 loop of MSI1 relative to the NMR structure, the number of native contacts between MSI1 and Numb RNA (*N*_*contacts*_), the radius of gyration (*R*_*g*_) and end-to-end distance of the Numb RNA were selected as reaction coordinates. Root-mean-square fluctuations (RMSFs) were calculated for the protein residues and RNA nucleotides, averaged over the GaMD production simulations and color coded for schematic representation of each system. Since only the Sim1 and Sim2 GaMD trajectories successfully captured complete binding of the Numb RNA to MSI1, these trajectories were used separately for structural clustering to identify the RNA binding pathways using the hierarchical agglomerative algorithm in CPPTRAJ^57^. The RMSD cutoff was set to 3.0 Å for the core RNA backbone to form a cluster.

The PyReweighting^52^ toolkit was applied to reweight GaMD simulations to recover the original free energy or PMF profiles of the two MSI1-Numb systems. All GaMD production simulations from the Unbound state (4,800 ns in total) and Bound state (1,500 ns in total) were combined for calculating the PMF profiles, respectively. A bin size of 1.0 Å was used for the core RNA backbone RMSD, the MSI1 β2-β3 loop backbone RMSD, the Numb *R*_*g*_ and the end-to-end distance of Numb. A bin size of 100 was used for *N*_*contacts*_. The cutoff was set to 500 frames for 2D PMF calculations.

## Results

### GaMD simulations captured complete binding of the Numb RNA to the MSI1 protein

Three independent 500 ns and four independent 1,200 ns GaMD simulations were performed on the MSI1-Numb system started from the Bound and Unbound states (**Table 1**). The GaMD simulations started from the Bound state recorded average and SD of the boost potential as 14.08 kcal/mol and 3.79 kcal/mol, respectively (**Table 1**). The GaMD simulations started from the Unbound state showed similar average and SD of boost potential with 11.53 kcal/mol and 3.36 kcal/mol, respectively (**Table 1**). The Bound MSI1-Numb complex was found to maintain the NMR structure with <3 Å RMSD of the core RNA backbone during most of the GaMD simulations (**Fig. 1A**). In the GaMD simulations started from the Unbound state, the core RNA backbone RMSD relative to the NMR structure decreased to minimum values 0.70 Å and 1.78 Å in Sim1 and Sim2, respectively, suggesting that complete binding of the Numb RNA from free diffusion in the solvent to the MSI1 target site was successfully captured (**Fig. 1b**). Spontaneous binding of RNA was observed in the Sim1 after ~100 ns with the RNA RMSD decreased to ~2.50 Å relative to the NMR structure (**Fig. 1B**). In Sim2, the Numb RNA bound to MSI1 during ~1,010-1,130 ns and then dissociated to the solvent (**Fig. 1B**). The RNA binding events captured in the present GaMD simulations allowed us to characterize the dynamic interactions between the MSI1 protein and Numb RNA.

**Figure 1.**
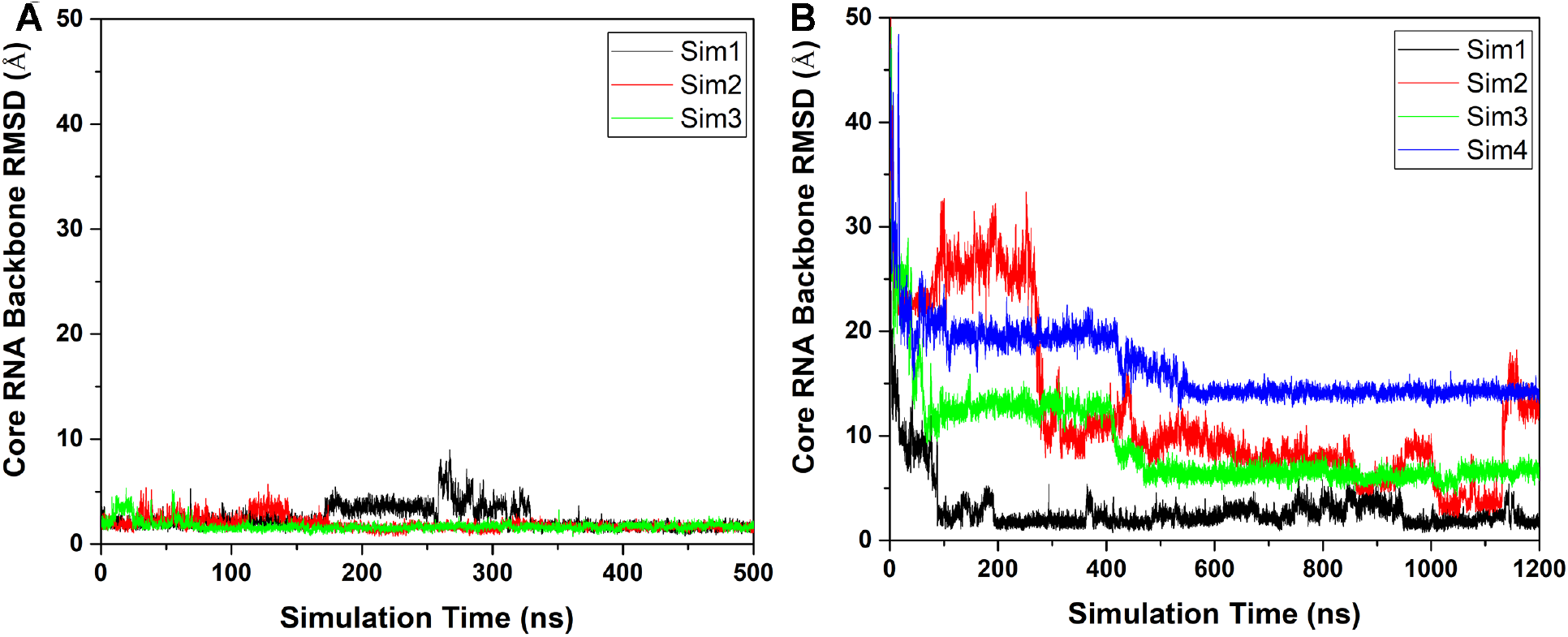
Time courses of the backbone RMSDs of core RNA (central three nucleotides GUA of the Numb RNA) relative to the NMR structure (PDB: 2RS2) are calculated from the GaMD simulations started from the (**A**) Bound and (**B**) Unbound states of the MSI1-Numb system.

### Variations of structural flexibility upon MSI1-Numb RNA binding

We analyzed structural flexibility of both the MSI1 protein and Numb RNA in the GaMD simulations. During GaMD simulations started from the Bound NMR structure, the MSI1 protein underwent small fluctuations except the loop connecting β2 and β3 strands (the β2-β3 loop) and the C terminus (**Fig. 2A**). The fifth nucleotide in the Numb RNA exhibited significantly higher flexibility than the other nucleotides, especially the central three ones UAG (denoted as the core RNA). This suggested that interactions between the core RNA and the MSI1 were strong. Thus, the core RNA might play an important role in the interactions between the MSI1 protein and Numb RNA. In compared to the GaMD simulations started from the Bound state, the Numb RNA exhibited much higher flexibility in the simulations started from the Unbound state. For the MSI1 protein, both the β2-β3 loop and C-terminus exhibited higher flexibility (**Fig. 2B**). These motifs were suggested to be important for binding of the Numb RNA^11^ and small molecules^14^ to the MSI1 protein.

**Figure 2.**
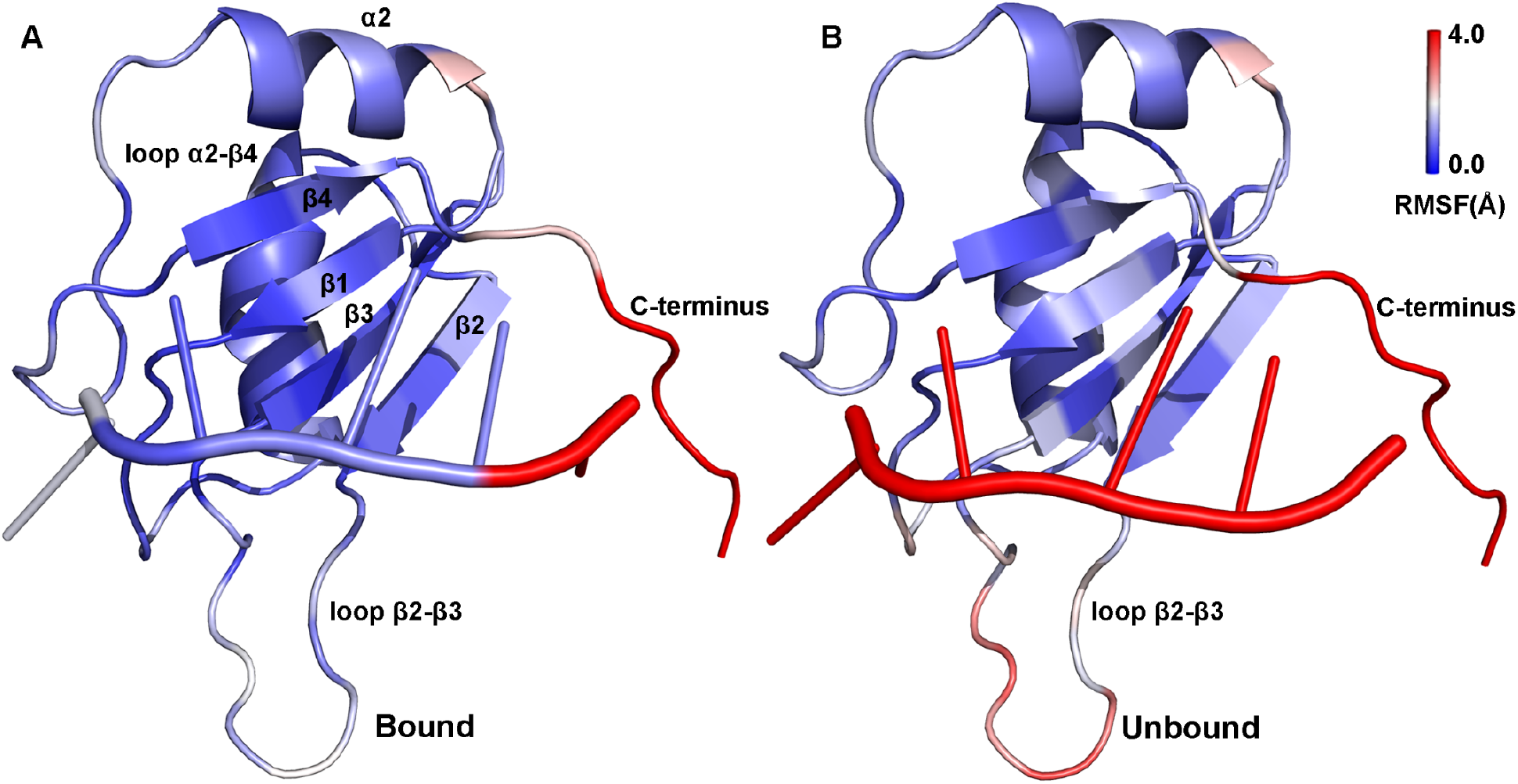
Structural flexibility of MSI1-RNA obtained from GaMD simulations: root-mean-square fluctuations (RMSFs) of the MSI1-RNA complex when the Numb RNA was initially placed in the (**A**) Bound state or (**B**) Unbound state. A color scale of 0.0 Å (blue) to 4.0 Å (red) is used.

### Free energy profiles of RNA binding to the MSI1 protein

Free energy profiles were calculated from the GaMD simulations using the core RNA backbone RMSD relative to the NMR structure and the number of native contacts between MSI1 and Numb RNA (*N*_*contacts*_) as reaction coordinates. Only one low-energy minimum of the “Bound” conformation was identified from the GaMD simulations on the NMR structure, in which the core RNA backbone RMSD and *N*_*contacts*_ centered around (1.2 Å, 1600) (**Fig. 3A**). Five low-energy minima were identified from GaMD simulations started from the Unbound state, including the “Bound”, “Intermediate I1”, “Intermediate I2”, “Intermediate I3” and “Unbound” states, in which the core RNA backbone RMSD and *N*_*contacts*_ centered around (2.0 Å, 1500), (5.2 Å, 600), (12.5 Å, 200), (25.0 Å, 10) and (40 Å, 0), respectively (**Fig. 3B**). The intermediate I1, I2 and I3 conformational states are shown in **Figs. 4**. Remarkably, positively charged residues (Arg61 and Arg99) in the β2-β3 loop and C-terminal region of the MSI1 protein formed favorable salt-bridge and hydrogen bond interactions with the central nucleotide A106 of Numb RNA. In the intermediate I1 state, the Numb RNA formed interactions with both the β2-β3 loop and C terminus of MSI1, leading to large conformational changes of these two regions (**Fig. 4A**). Notably, the sidechain of residue Arg99 in the C terminus of the MSI1 protein formed three hydrogen bonds with the sidechain of nucleotide A106 in the Numb RNA (**Fig. 4A**). In the intermediate I2 state, the sidechain of residue Arg61 in MSI1 could flip out to the solvent, forming a hydrogen bond and a salt-bridge with the sidechain and backbone (oxygen atom in the phosphate group) of the nucleotide A106 in Numb RNA, respectively (**Fig. 4B**). In the intermediate I3 state, Arg99 in the C terminus of MSI1 formed a hydrogen bond and a salt-bridge with sidechain and backbone of the nucleotide A106 in Numb RNA, respectively, for which a large conformational change of the protein C terminus was observed (**Fig. 4C**). Thus, these hydrogen bonds and salt-bridge interactions played a significant role in the recognition and binding of the Numb RNA to MSI1 protein. Since the charged residues Arg61 and Arg99 were identified to interact with the Numb RNA via hydrogen bond and long-range salt-bridge interactions in the intermediate states, we assumed that the Numb RNA binding to the MSI1 protein was mainly mediated by electrostatic interactions.

**Figure 3.**
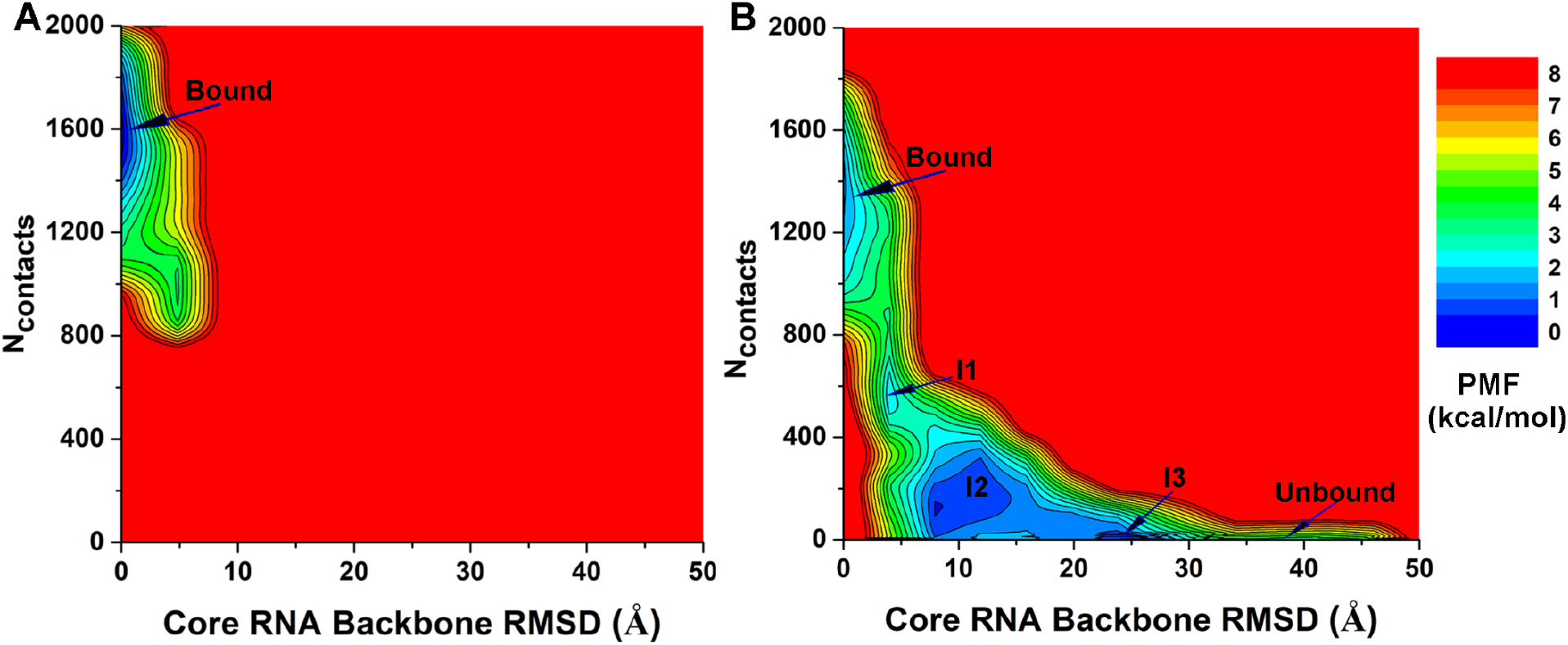
2D potential of mean force (PMF) free energy profiles of the core RNA backbone RMSD relative to the NMR structure (PDB: 2RS2) and number of native contacts between MSI1 and Numb RNA are calculated from GaMD simulations started from the (**A**) Bound and (**B**) Unbound states of the MSI1-Numb system.

**Figure 4.**
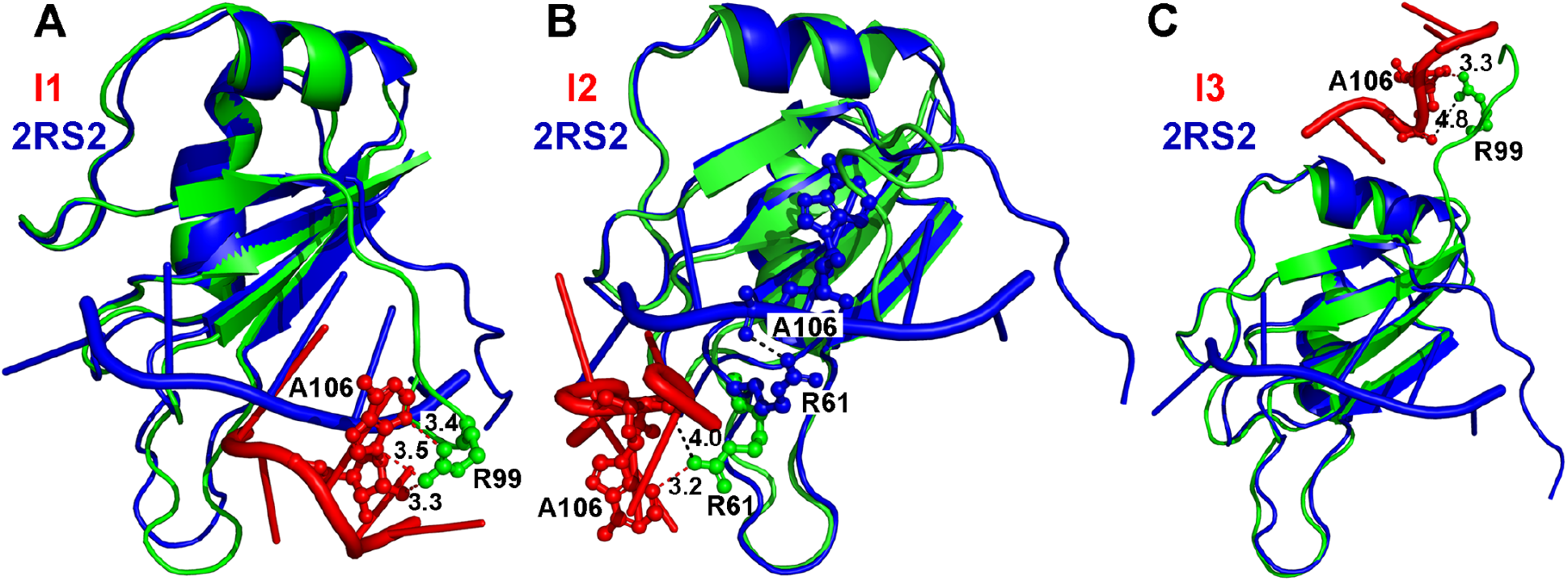
Low-energy intermediate conformational states (I1, I2 and I3) as identified from the 2D PMF profile of the MSI1-RNA simulation system started from the Unbound state. The MSI1 protein and Numb RNA are shown in green and red, respectively. The NMR structure of the MSI1-Numb complex is shown in blue for comparison. MSI1 protein residues Arg99 and Arg61, and nucleotide A106 of Numb RNA are highlighted in balls and sticks. The hydrogen bonds between the side-chain of residues in the MSI1 protein and Numb RNA are shown in red. The salt-bridge interactions between the side-chain of residues in the MSI1 protein and backbone (oxygen atom) of the Numb RNA are shown in black.

As described above, binding of the Numb RNA induced higher flexibility of the MSI1 β2-β3 loop (**Fig. 1B**) and large conformational change of the same region was observed in the intermediate I1 state (**Fig. 4A**). Therefore, the MSI1 β2-β3 loop backbone RMSD and core Numb RNA backbone RMSD relative to the experimental structure were used as reaction coordinates to further compute 2D free energy profiles (**Figs. 5)**. The MSI1 β2-β3 loop was highly flexible, sampling a large conformational space with the backbone RMSD ranging from ~0 Å to ~8.0 Å (**Fig. 5B**). This loop sampled two distinct low-energy conformations, including the “Closed” (bound) (RMSD < 1 Å) and “Open” (free) states (RMSD ~3-5 Å) (**Fig. 5B**). Compared to the “Open” state, the MSI1 β2-β3 loop moved closer to the core domain in the “Closed” state (**Fig. 4B**). Five low-energy states were identified from GaMD simulations starting with the Unbound state, including the “Unbound/Open”, “Intermediate I3/Open”, “Intermediate I2/Open”, “Intermediate I1/Closed” and “Bound/Open” states, in which the MSI1 β2-β3 loop backbone RMSD and core RNA backbone RMSD were located around (1.0 Å, 40 Å), (1.2 Å, 25.0 Å), (1.1 Å, 12.5 Å), (4.5 Å, 7.5 Å) and (1.5 Å, 2.0 Å), respectively (**Fig. 5B**). The Numb RNA and MSI1 β2-β3 loop accommodated to each other to form the final bound conformation (**Fig. 5B**), suggesting an “induced fit” mechanism in the MSI1-Numb RNA interaction.

**Figure 5.**
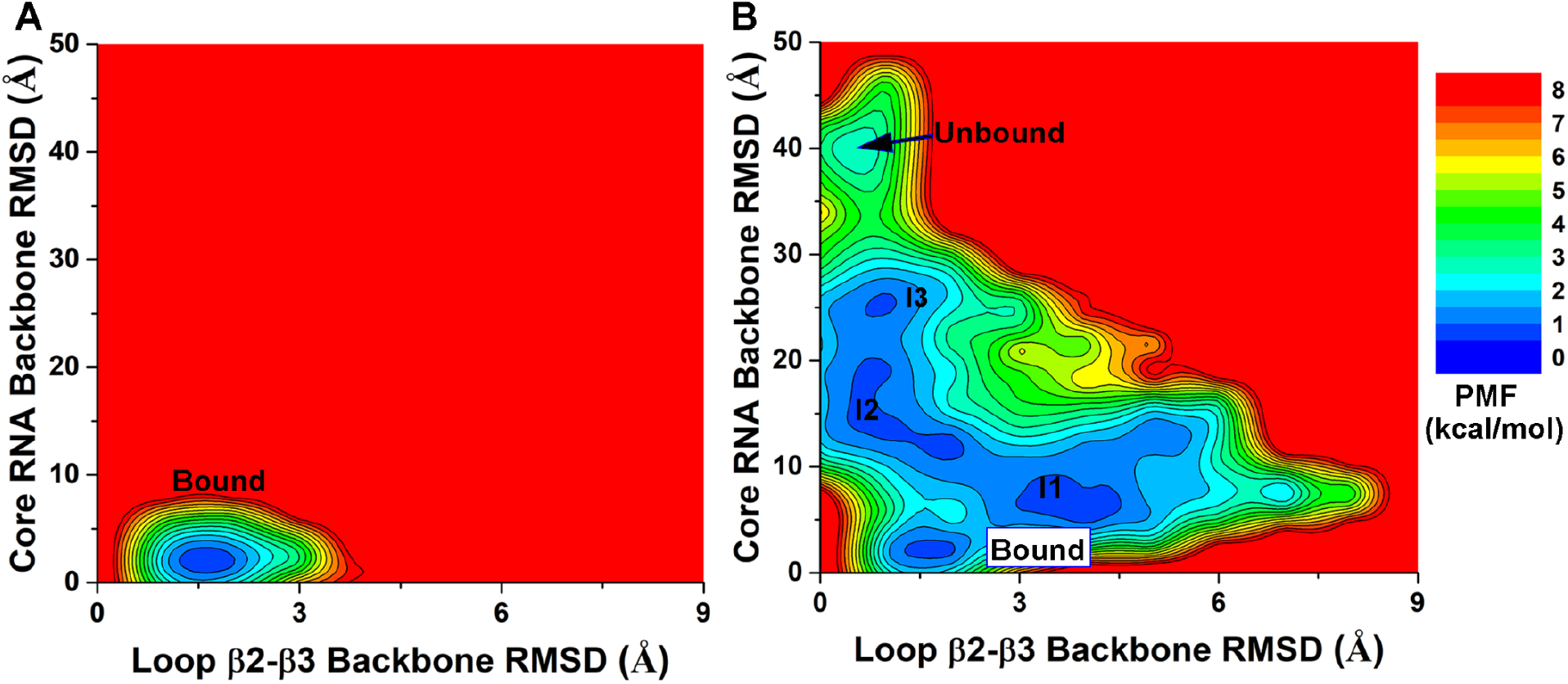
2D PMF profiles of the MSI1 β2-β3 loop backbone RMSD and core RNA backbone RMSD relative to the NMR structure (PDB: 2RS2) are calculated from GaMD simulations started from the (**A**) Bound and (**B**) Unbound states of the MSI1-Numb system.

### Pathways of RNA binding to the MSI1 protein

Next, structural clustering was performed separately on the GaMD trajectories of Sim1 (movie S1 in Supporting Information) and Sim2 (movie S2 in Supporting Information) to identify the representative binding pathways of the Numb RNA to the MSI1 protein. The structural clusters were reweighted to obtain their original free energy values, which ranged from 0.0 kcal/mol to ~4.5 kcal/mol. The top reweighted clusters with PMF ≤2.0 kcal/mol were selected to represent the pathways of the Numb RNA binding to MSI1 (**Fig. 6)**. In Sim1, the Numb RNA bound to MSI1 via interactions with the protein C terminus (**Fig. 6A**). In Sim2, both the β2-β3 loop and C terminus of MSI1 contributed important interactions with the Numb RNA during its binding to the protein target site (**Fig. 6B**). These findings again revealed important roles of the β2-β3 loop and C terminus of MSI1 in binding of the Numb RNA. This was consistent with the above RMSF analysis that higher flexibilities were observed in these two dynamic regions of MSI1 (**Fig. 2**).

**Figure 6.**
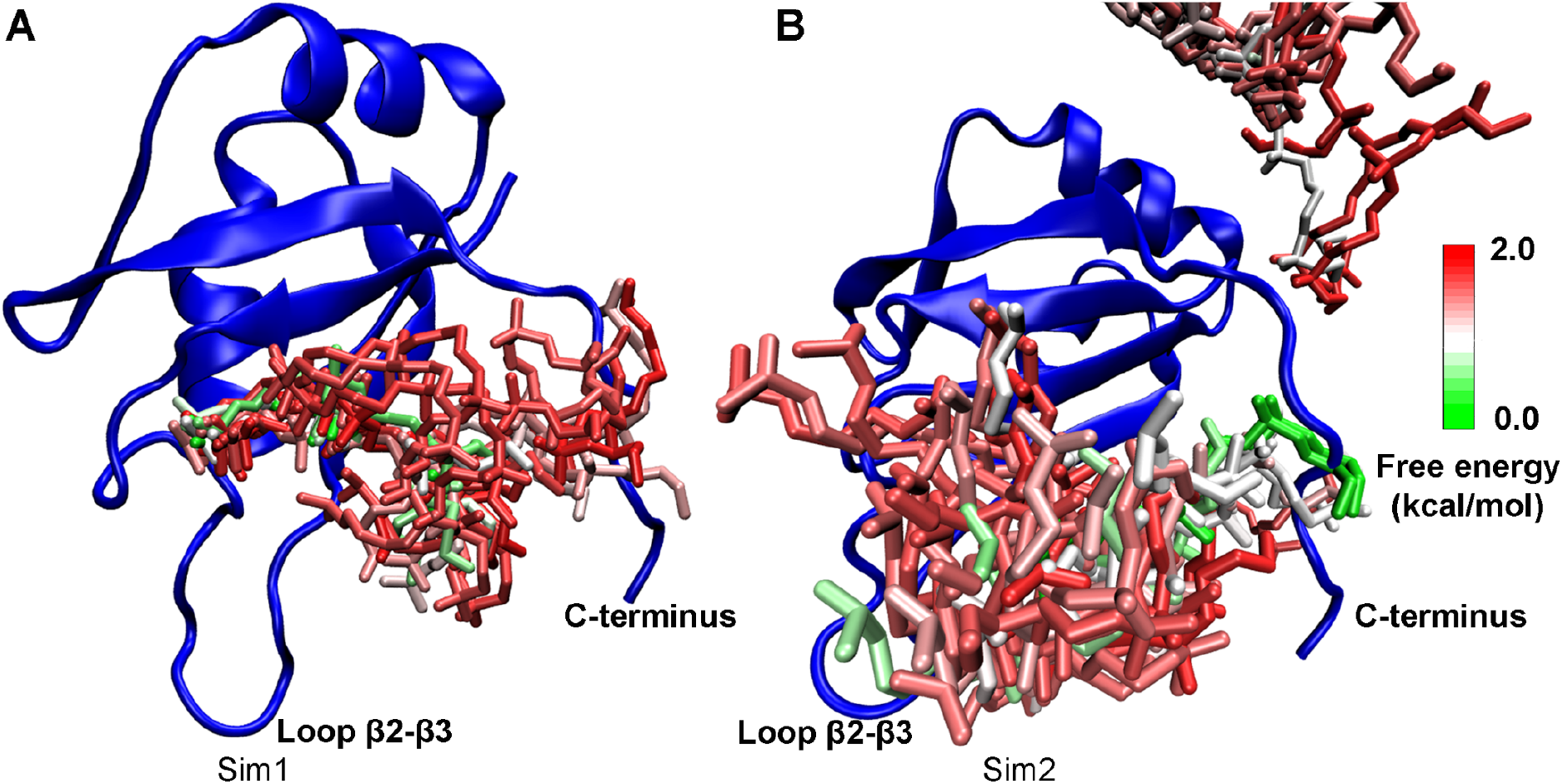
Binding pathways of the Numb RNA to the MSI1 protein revealed from GaMD simulations: **(A)**Starting from diffusion in the solvent, the Numb RNA bound to the target site of the MSI1 via intermediate conformations by interacting with the MSI1 C-terminus in the “Sim1” GaMD trajectory. (**B**) Starting from diffusion in the solvent, the Numb RNA bound to the target site of the MSI1 protein via intermediate conformations by interacting with the MSI1 β2-β3 loop and C-terminus in the “Sim2” GaMD trajectory. The MSI1 is shown in blue ribbons. The Numb structural clusters (sticks) are colored by the reweighted PMF free energy values in a green (0.0 kcal/mol)-white-red (2.0 kcal/mol) color scale.

### The Numb RNA bound to the MSI1 protein via an induced fit mechanism

In order to further explore the mechanism of RNA binding to the MSI1 protein, we computed free energy profiles to characterize conformational changes of the Numb RNA upon binding to MSI1. In this regard, the radius of gyration (*R*_*g*_) and the end-to-end distance of Numb were calculated to monitor its possible conformational changes. We used the *R*_*g*_ and end-to-end distance of the Numb RNA and the core RNA backbone RMSDs as reaction coordinates to calculate 2D PMF profiles (**Figs. 7A-7D**). Notably, the Numb RNA sampled a large conformational space during binding to the MSI1 protein in the GaMD simulations started from the Unbound state (**Figs. 7B**). From the reweighted 2D PMF profiles, we identified a similar “Bound” low-energy well in simulations started from both the Bound and Unbound states, for which the Numb RNA *R*_*g*_ and core RNA backbone RMSD centered around (2.0 Å, 8.5 Å) and (2.5 Å, 8.2 Å), respectively (**Fig. 7A-7B**). Another four low-energy states were identified in GaMD simulations started from Unbound conformation, including the “Unbound”, “Intermediate I1”, “Intermediate I2” and “Intermediate I3”, for which the core RNA backbone RMSD and *R*_*g*_ of Numb centered around (40.0 Å, 6.2 Å), (5.7 Å, 6.5 Å), (5.7 Å, 12.5 Å), respectively (**Fig. 7B**).

**Figure 7.**
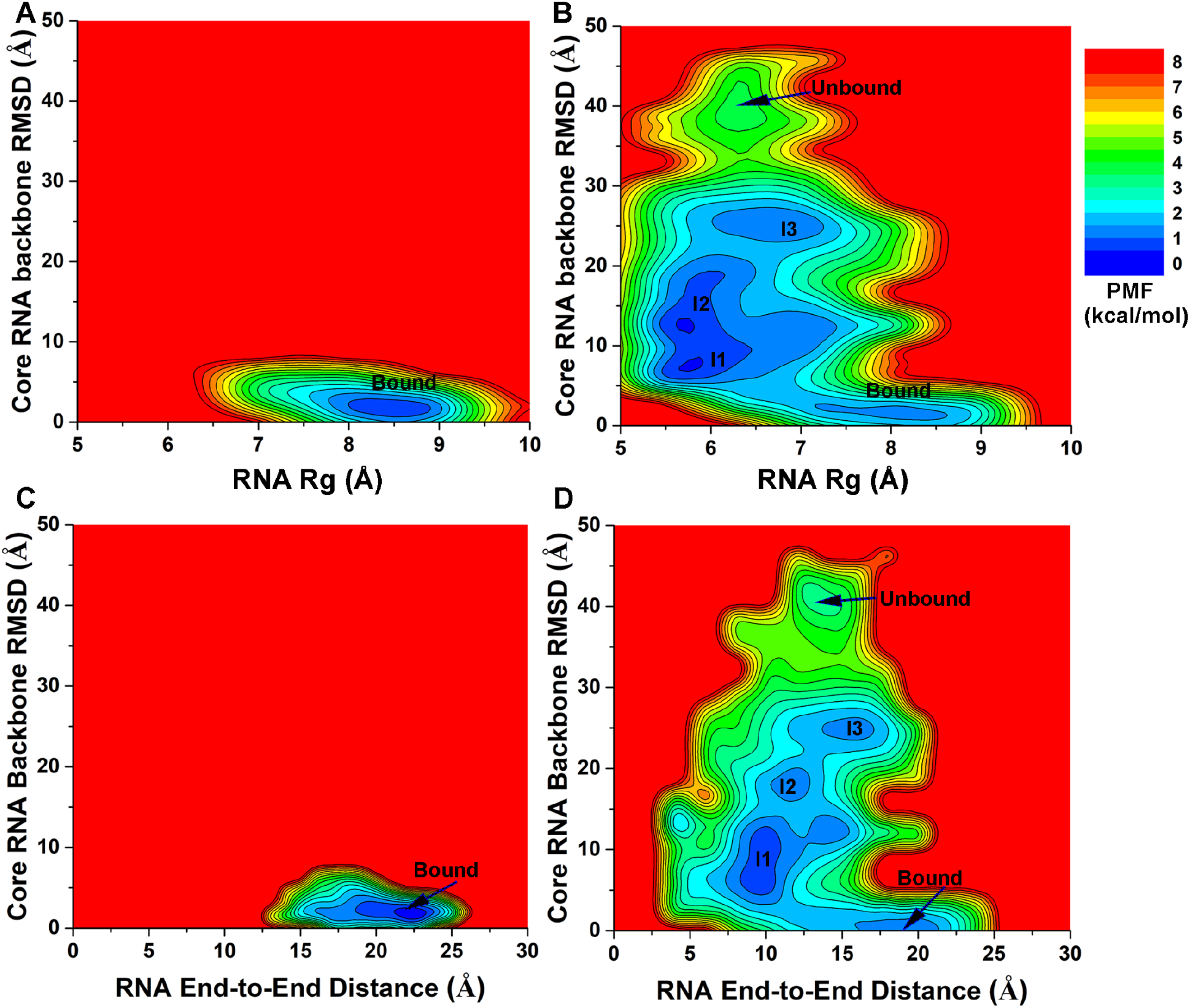
(**A-B**) 2D PMF profiles of the radius of gyration (*R*_*g*_) of the Numb RNA and core RNA backbone RMSD relative to the NMR structure (PDB: 2RS2) are calculated from GaMD simulations started from the (**A**) Bound and (**B**) Unbound states of the MSI1-Numb system. (**C-D**) 2D PMF profiles of the end-to-end distance of the Numb RNA and core RNA backbone RMSD relative to the NMR structure (PDB: 2RS2) are calculated from GaMD simulations started from the (**C**) Bound and (**D**) Unbound states of the MSI1-Numb system.

Furthermore, we calculated 2D PMF profiles regarding the core RNA backbone RMSD and the end-to-end distance of Numb RNA (**Figs. 7C-7D**). Only one low-energy state (“Bound”) was identified in the 2D PMF profile calculated from the GaMD simulations of the bound NMR structure, in which the Numb RNA adopted primarily the “Extended” conformation (**Fig. 7C**). The Numb end-to-end distance and core RNA backbone RMSD centered around (22.5 Å, 1.8 Å) (**Fig. 7C**). In contrast, five distinct low-energy states were identified from the 2D PMF profile in the GaMD simulations started from the Unbound conformation, including the “Unbound”, “Intermediate I1”, “Intermediate I2”, “Intermediate I3” and “Bound”, in which the Numb RNA adopted primarily the “Curled”, “Curled”, “Curled”, “Curled”, and “Extended” conformations, respectively (**Figs. 7D** and **4**). The end-to-end distance of Numb RNA and core RNA backbone RMSD centered around (12.5 Å, 40.0 Å) in the “Unbound/Curled” state, (14.0 Å, 25.0 Å) in the “Intermediate I3/Curled” state, (18.5 Å, 10.5 Å) in the “Intermediate I2/Curled” state, (9.5 Å ,7.5 Å) in the “Intermediate I1/Curled” state and finally (20.0 Å, 2.5 Å) in the “Bound/Extended” state (**Fig. 7D**).

In comparison, the Numb RNA sampled a larger conformational space with a wider range of *R*_*g*_ or end-to-end distance in the “Bound” state than in the “Unbound” and even the three Intermediate conformational states (**Figs. 7B** and **7D**). Therefore, binding of the Numb RNA to the MSI1 protein involved largely induced fit.

## Discussion

In this study, we have applied all-atom GaMD simulations to investigate dynamic interactions between the Numb RNA and MSI1 protein. The GaMD simulations unprecedentedly captured spontaneous and highly accurate binding of the Numb RNA from bulk solvent to the MSI1 protein with <2 Å RMSD in the core RNA backbone compared with the experimental structure. Proper energetic reweighting of the GaMD simulations allowed us to calculate PMF free energy profiles to characterize the MSI1-Numb binding process.

Important low-energy conformational states were identified from the GaMD derived free energy profiles. Particularly, in the I1, I2 and I3 intermediate states, a significant role of the β2-β3 loop and C terminus of the MSI1 protein was revealed in the recognition and binding of the Numb RNA (**Fig. 4**). The charged residues Arg61 and Arg99 of MSI1 that formed hydrogen bond and salt-bridge interactions with the Numb RNA were observed in the GaMD simulations, supporting important roles of electrostatic interactions in forming the “Intermediate” and “Bound” states of the MSI1-Numb RNA complex. The important role of Arg99 was consistent with the result that the mutation of the same residue in its homology MSI2 (Arg100Ala) decreased its binding affinity of Numb^10^. The salt-bridge between Arg61 (MSI1 protein) and A106 (Numb RNA) was also observed in the bound NMR structure (**Figs. 4B**). In contrast, our GaMD simulations here captured this interaction during the binding process of the Numb RNA, supporting the important role of electrostatic interactions in forming both the “Intermediate” and “Bound” states. Furthermore, Arg61 was characterized as important residue for the inhibitor binding as the MSI1 Arg61Glu mutant decreased ~5 fold for the binding of inhibitor^14^. Therefore, GaMD simulations showed that long-range electrostatic interactions played an important role in the Numb RNA binding to the MSI1 protein and identified two critical residues (Arg61 and Arg99) in the binding process. Additionally, RMSF analysis suggested that the binding between the core RNA and the MSI1 protein was highly stable in the GaMD simulations starting from the Bound state, suggesting that the core RNA might play an important role in the interactions between the MSI1 protein and the Numb RNA. This result is consistent with the result obtained by Zearfoss et al.^59^ that the central core RNA (UAG) forms the core MSI recognition element and makes major contributions to binding affinity.

Conformation selection^60–62^ and induce fit^63–64^ are two common models for describing biomolecular recognition. In this context, our GaMD simulations have revealed that binding of the Numb RNA to the MSI1 protein involved predominantly an induced fit mechanism, in which both the RNA and protein (especially the β2-β3 loop) underwent significant conformational changes during binding. This is consistent with previous studies of other protein-RNA interactions^64–67^, including ribosomal protein S15-rRNA and U1A-RNA complexes. A major conformational change of the rRNA was found upon binding to the S15 protein through comparison of the free and bound structures of S15 and rRNA, suggesting induced fit of the protein and RNA^64^. MD simulations combined with available structure analysis also indicated that binding of the U1A protein and RNA followed an induced fit mechanism^67^. For the U1A protein, MD simulations indicated that induced fit upon binding involved a non-native thermodynamic substate, in which the structure is preorganized for binding. In contrast, induced fit of the RNA involved a distortion of the native structure to an unstable form in solution.

It is worth noting that the presented free energy profiles were not converged since only two binding events were observed in the GaMD simulations. More binding and unbinding events would still need to be simulated in order to quantitatively characterize RNA binding to the MSI1 protein and calculate the binding free energies and kinetic rates. In this regard, our recently developed selective GaMD algorithms^68–69^ could be useful to address the challenge. In particular, microsecond simulations using the peptide GaMD (Pep-GaMD) method^69^, which works by selectively boosting the peptide potential energy, have been demonstrated to capture repetitive binding and unbinding of highly flexible peptides to the target protein^69^. Apart from enhanced conformational sampling, accurate force fields are also needed especially for the RNA^70–73^ in order to simulate repetitive RNA dissociation and binding to RBPs. Even the force field works well individually for the protein and RNA, combination of protein and RNA force fields in the MD simulations could introduce additional challenges.^18, 20, 70^ Nevertheless, we have observed two complete binding events of the Numb RNA to the MSI1 protein with our current force field settings and GaMD simulations, which will guide our future simulation studies of RBPs-RNA interactions.

## Conclusions

In summary, all-atom GaMD simulations with unconstrained enhanced sampling and free energy calculations have provided important insights into the mechanism of the Numb RNA binding to the MSI1 protein. For future studies, the effects of small molecule binding in the MSI1-Numb interactions still need to be determined and our simulation findings await validation in the wet-lab experiments. Further studies are planned to simulate both dissociation and binding of RNA to the RBPs and accurately predict the thermodynamics and kinetics of protein−RNA interactions. These efforts are expected to greatly facilitate rational drug design targeting the MSI1 and other RBPs.

## Author Contributions

Y.M. designed research. J.W. performed the research. J.W., L. L, X.W., L. X and Y.M. analyzed the data, and J.W. and Y.M. wrote the paper.

## Conflict of Interest Statement

The authors declare that the research was conducted in the absence of any commercial or financial relationships that could be construed as a potential conflict of interest.

## Acknowledgements

This work was supported in part by the National Institutes of Health (R01GM132572) and the startup funding in the College of Liberal Arts and Sciences at the University of Kansas (to Y.M.), NIH R01 CA191785 and KU Cancer Center Pilot Project Award (to L.X.). This work used supercomputing resources with allocation Award No. TG-MCB180049 through the Extreme Science and Engineering Discovery Environment (XSEDE), which is supported by the National Science Foundation, Grant No. ACI-1548562, and Project No. M2874 through the National Energy Research Scientific Computing Center (NERSC), which is a U.S. Department of Energy Office of Science User Facility operated under Contract No. DE-AC02-05CH11231, and the Research Computing Cluster at the University of Kansas.

## Supplementary Material

The Supporting Information is available free of charge on the ACS Publications website. Movies S1 and S2 showing GaMD trajectories of Sim1 and Sim2 starting from the Unbound state, respectively.

## Table of content

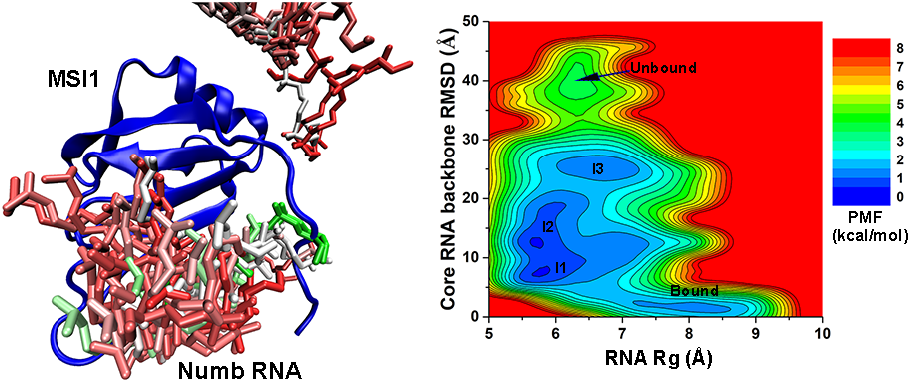
Abstract Graphic. All-atom simulations using a robust Gaussian accelerated molecular dynamics (GaMD) method successfully captured spontaneous binding of the Numb RNA to the Musashi-1 (MSI1) protein, which involved largely induced fit in both the RNA and protein. GaMD simulations also identified important low-energy intermediate conformational states of RNA binding. The simulation findings are expected to facilitate rational structure-based drug design targeting MSI1 and other RNA-binding proteins.

## Notes

### Competing Interest Statement

The authors have declared no competing interest.

